# Halophilic microbial community compositional shift after a rare rainfall in the Atacama Desert

**DOI:** 10.1101/442525

**Authors:** Gherman Uritskiy, Samantha Getsin, Adam Munn, Benito Gomez-Silva, Alfonso Davila, Brian Glass, James Taylor, Jocelyne DiRuggiero

## Abstract

Understanding the mechanisms underlying microbial resistance and resilience to perturbations is essential to predict the impact of climate change on Earth’s ecosystems. However, the resilience and adaptation mechanisms of microbial communities to natural perturbations remain relatively unexplored, particularly in extreme environments. The response of an extremophile community inhabiting halite (salt rocks) in the Atacama Desert to a catastrophic rainfall provided the opportunity to characterize and de-convolute the temporal response of a highly specialized community to a major disturbance. With shotgun metagenomic sequencing, we investigated the halite microbiome taxonomic composition and functional potential over a 4-year longitudinal study, uncovering the dynamics of the initial response and of the recovery of the community after a rainfall event. The observed changes can be recapitulated by two general modes of community shifts – a rapid *Type 1* shift and a more gradual *Type 2* adjustment. In the initial response, the community entered an unstable intermediate state after stochastic niche re-colonization, resulting in broad predicted protein adaptations to increased water availability. In contrast, during recovery, the community returned to its former functional potential by a gradual shift in abundances of the newly acquired taxa. The general characterization and proposed quantitation of these two modes of community response could potentially be applied to other ecosystems, providing a theoretical framework for prediction of taxonomic and functional flux following environmental changes.

## INTRODUCTION

Microbial communities are essential to the functioning and evolution of our planet and their dynamics greatly affect ecosystems processing (1). Their taxonomic and functional diversity allow microbial communities to adapt to a wide range of conditions and to respond rapidly to environmental changes (2, 3). Resilience – the ability of a community to recover from perturbations – is of particular interest, especially in the context of global climate change, as extreme weather events are becoming more frequent (1). Understanding adaptation strategies for microbial resilience is therefore critical to gain insights into microbial evolution and diversification and to better understand the dynamics of translationally relevant microbiomes following stress.

Previous studies have shown that acute disturbances can push a community’s taxonomic structure toward alternative equilibrium states, while retaining the preexisting functional potential (4). Such changes have been observed in soil, aquatic, engineered, and human-associated ecosystems where experimental perturbations caused the community taxonomic composition to shift with relatively minor changes to the overall functional potential of the community (1, 2, 5, 6). Functional redundancy has been proposed as a mechanism to support functional stability following perturbation (7), however several studies have shown that major taxonomic changes can also result in important changes to the functional potential of gut communities (8, 9).

Transitions between alternative taxonomic states have been postulated to occur via an intermediate dis-equilibrium state, during which a perturbation produces drastically different environmental stressors, causing the community to radically reshape in composition (1, 4). This has been observed with antibiotic treatment that can lead to mass death events. The resulting re-structuring of the gut microbiome is major with long-lasting changes even after the former conditions are re-established (6, 10). However, little is known about the response dynamics to acute perturbations and in particularly the mechanisms that push a community’s taxonomic and functional structure in and out of an intermediate state. Additionally, the response and recovery of natural communities following environmental disasters, rather than manipulative experiments, remain largely unexplored mechanistically because of the difficulty in avoiding multiple compounding environmental factors (11, 12). These gaps in the understanding of microbial community behavior limits our ability to effectively model and predict the responses of microbiomes to major perturbations, such as those resulting from climate change and natural or man-made ecological disasters.

To address this knowledge gap, and to build a conceptual model for modeling microbial community responses to extreme stress, we examined the temporal dynamics in response to a disastrous climate perturbation of a unique microbial ecosystem found in the Atacama Desert, Chile. The hyper-arid core of the Atacama Desert is one of the harshest environments on Earth, with an average annual precipitation of less than 1mm and some of the highest ultraviolet (UV) and solar radiation on the planet (13, 14). Despite this, microbial communities have evolved strategies to survive and grow within various mineral substrates of the desert (15). One such community inhabits halite nodules that are natural porous salt rocks found exclusively in evaporitic salt basins of the Atacama Desert, including the Salar Grande basin (16, 17) (Fig. S1). In this community, the majority of the biomass is constituted of salt-in strategists Halobacteria (a major class of archaea) and Bacteroidete*s* (17, 18) – two taxonomically diverse groups of extreme halophiles that accumulate potassium ions to match the external osmotic pressure from sodium ions (17, 19, 20). This adaptation allows them to survive in extremely high-salt environment, but restricts their fitness to a narrow range of external salt concentration (21, 22). As such, these highly specialized communities are more vulnerable to change compared to habitat generalists, particularly to sudden changes in external osmotic pressure.

Encased in salt rocks, halite communities have very limited nutrient input beyond atmospheric gasses, and obtain water almost exclusively from deliquescence, the ability of sodium chloride to produce concentrated brine when atmospheric relative humidity rises above 75% (23). Primary production is the major source of organic carbon in the community and is carried out by *Cyanobacteria* and, to a lesser extent, by a unique alga *(17)*. Each halite nodule represents a near-closed miniature ecosystem and thus can be treated as true independent biological replicates in longitudinal studies, allowing community changes to be tracked without external factors compounding the results. Combined with their sensitivity to changing osmotic conditions and slow growth rates, this makes halite microbiomes ideal for studying temporal dynamics of microbial communities and their ability to adapt to major environmental changes.

In 2015, Northern Atacama received its first major rain in 13 years (14). In particular, a weather station located 40 km North-West of our sampling site (Diego Aracena Airport *SCDA*) recorded significant rainfalls of 4.1mm (August 9^th^, 2015) and 20.1mm (November 20^th^, 2015) (24). The previous notable precipitation in the area occurred in 2002 (4.1mm) (25). Such rain events have been observed to be devastating to the specialized hyper-arid microbiomes of the Atacama Desert (26), particularly in communities adapted to survive in saturated salt conditions, such as those found in halite nodules. Our longitudinal study over 4 years not only captured the microbiome’s short-term adaptations to this major natural disaster, but also its recovery in the subsequent years, revealing two strikingly different community response mechanisms.

## MATERIALS AND METHODS

### Longitudinal sampling strategy and sequencing approach

To investigate the temporal dynamics of halite microbiomes, samples of halite nodules from two sites at Salar Grande (Fig. S1), a salar in the Northern part of the Atacama Desert (16), were harvested at regular intervals from 2014 to 2017, capturing the rare rain events that occurred in 2015 throughout the desert (14). The main sampling site (Site 1) was revisited four times during the study – twice before the rain (Sep 2014, Jun 2015), and twice after the rain – 3 months (Feb 2016) and 15 months (Feb 2017) after (Table S1). For each time-point, 5 biological replicates were sequenced with whole-metagenomic (WMG) shotgun sequencing to investigate the functional potential and taxonomic structure of the communities over time, yielding a total of 70,689,467 paired-end reads (150bp paired-end, insert size 277±217bp). Additionally, 9-12 biological replicates were collected for ribosomal amplicons (16S rRNA gene) sequencing and were used for taxonomic profiling of the microbiomes; this yielded 535,233 paired-end reads (250bp paired-end, insert size 419±7bp). A nearby site (Site 2) was also sampled after the rain at a higher temporal resolution (Feb 2016, July 2016, Oct 2016, and Feb 2017), with 5-13 replicates per time point. The 16S rRNA gene amplicons from samples at this site were also sequenced, yielding 357,325 paired end 250bp reads (insert size 419±4bp).

### Climate data acquisition

Climate history data was obtained from the Weather Underground weather reporting service by selecting “Monthly History” in the data browser (24). Weather data collected at the Diego Aracena International Airport (code SCDA) was manually downloaded for dates from the duration of the study (Jan 2014 – Mar 2017). The minimum and maximum temperature and relative humidity, as well as total precipitation data from each day were plotted against time. The raw unedited data and analysis scripts can be found at https://github.com/ursky/timeline_paper.

### Sample collection and DNA extraction

To investigate the effect of the rain on different locations, halite nodules were harvested from three sites in Salar Grande (Table 1). At each site, halite nodules were harvested within a 50m^2^ area. At the S1 location, 14-24 replicates were collected yearly over the course of 4 years for the main analysis in this work comparing pre- and post-rain samples with both shotgun and amplicon sequencing. At the S2 location, 5-13 replicates were collected from 4 time points in the year following the rain to validate the post-rain community recovery with amplicon sequencing. Finally, shotgun sequencing of samples from the S3 location were used to improve the binning results from S1, but were not used for the longitudinal analysis of this work because too few time points and replicates were collected (see Table S1 for details on sampling sites and replication). Halite nodules were collected as previously described (16) and ground into a powder, pooling material from 1-3 larger nodules until sufficient amount was collected, and stored in dark in dry conditions until DNA extraction in the lab. Genomic DNA was extracted as previously described (16, 17) with the DNAeasy PowerSoil DNA extraction kit (QIAGEN).

### 16S rRNA gene amplicon library preparation and sequencing

The communities’ 16S rRNA gene was amplified with a 2-step amplification and barcoding PCR strategy as previously described (16) by amplifying the hypervariable V3-V4 region with 515F and 926R primers (27). PCR was done with the Phusion High-Fidelity PCR kit (New England BioLabs) with 40ng of gDNA. Barcoded samples were quantified with the Qubit dsDNA HS Assay Kit (Invitrogen), pooled and sequenced on the Illumina MiSeq platform with 250 bp paired-end reads at the Johns Hopkins Genetic Resources Core Facility (GRCF).

### Shotgun metagenomic library preparation

Whole-genome metagenomic sequencing libraries were prepared using the KAPA HyperPlus kit (Roche). The fragmentation was performed with 5ng of input gDNA for 6 minutes to achieve size peaks of 800bp. Library amplification was done with dual-index primers for a total of 7 cycles, and the product library was cleaned 3 times with XP AMPure Beads (New England BioLabs) to remove short fragments and primers (bead ratios 1X and 0.6X, keep beads) and long fragments (0.4X bead ratio, discard beads). Other steps followed the manufacturer’s recommendations. The final barcoded libraries were quantified with Qubit dsDNA HS kit, inspected on a dsDNA HS Bioanalyzer, pooled to equal molarity, and sequenced with paired 150bp reads on the HiSeq 2000 platform at GRCF.

### 16S rRNA gene amplicon sequence analysis

The de-multiplexed and quality trimmed 16S rRNA gene amplicon reads from the MiSeq sequencer were processed with MacQIIME v1.9.1 (28). Samples from site 1 and 2 were processed separately. The reads were clustered into OTUs at a 97% similarity cutoff with the pick_open_reference_otus.py function (with --suppress_step4 option), using the SILVA 123 database (29) release as reference and USEARCH v6.1.554 (30). The OTUs were filtered with filter_otus_from_otu_table.py (-n 2 option), resulting in a total of 472 OTUs for site 1 and 329 OTUs for site 2 (Data S1). The taxonomic composition of the samples was visualized with summarize_taxa_through_plots.py (default options; Data S1). The beta diversity metrics of samples from the two sites were compared by normalizing the OTU tables with normalize_table.py (default options), and then running beta_diversity.py (-m unweighted_unifrac, weighted_unifrac). The sample dissimilarity matrices were visualized on PCoA plots with principal_coordinates.py (default parameters) and clustered heat maps with clustermap in Seaborn v0.8 (31) (method=‘average’, metric=‘correlation’). Group significance was determined with compare_categories.py (--method=permanova). Relative similarity between metadata categories (harvest dates) was calculated with the make_distance_boxplots.py statistical package, which summarized the distances between pairs of sample groups (from Weighted or Unweighted Unifrac dissimilarity matrices), and then performed a two-sided Student’s two-sample t-test to evaluate the significance of differences between the distances. Relative abundance of phyla and domain taxa were computed from the sum of abundances of OTUs with their respective taxonomy, and group significance calculated with a two-sided Student’s two-sample t-test. Detailed scripts for the entire analysis pipeline can be found at https://github.com/ursky/timeline_paper.

### Processing shotgun metagenomic sequence data

The de-multiplexed WMG sequencing reads were processed with the complete metaWRAP v0.8.2 pipeline (32) with recommended databases on a UNIX cluster with 48 cores and 1024GB of RAM available. Read trimming and human contamination removal was done by the metaWRAP Read_qc module (default parameters) on each separate sample. The taxonomic profiling was done on the trimmed reads with the metaWRAP Kraken module (33) (default parameters, standard KRAKEN database, 2017). The reads from all samples from the 3 sampling sites were individually assembled (for *pI* calculations) and co-assembled (for all other analysis) with the metaWRAP Assembly module (--use-metaspades option) (34). For improved assembly and binning of low-abundance organisms, reads from all samples were co-assembled, then binned with the metaWRAP Binning module (--maxbin2 --concoct --metabat2 options) while using all the available samples for differential coverage information. The resulting bins were then consolidated into a final bin set with metaWRAP’s Bin_refinement module (-c 70 -x 5 options; Data S2). The bins were then quantified by Salmon (35) with the Quant_bins module (default parameters). Contig read depth was estimated for each sample with the metaWRAP’s Quant_bins module, and the weighted contig abundance calculated by multiplying the contig’s depth by its length, and standardizing to the total contig abundance in each replicate. Detailed scripts for the entire analysis pipeline can be found at https://github.com/ursky/timeline_paper.

### Functional annotation

Gene prediction and functional annotation of the co-assembly was done with the JGI Integrated Microbial Genomes & Microbiomes (IMG) (36) annotation service. Gene relative abundances were taken as the average read depth of the contigs carrying those genes (estimated with Salmon (35). KEGG KO identifiers were linked to their respective functions using the KEGG BRITE pathway classification (37). KEGG pathway relative abundances were calculated as the sum of read depths of genes (estimated from the read depths of the contigs carrying them) classified to be part of the pathway. To test for changes in functional diversity, the total number of unique enzyme identifiers that had a combined coverage of 1, 2, 4, 8, 16, or 32 transcripts per million was calculated.

### Isoelectric point (*pI*) analysis

The average <u>*pI*</u> of gene pools were calculated from individual replicate metagenomic assemblies. Open reading frames (ORFs) were predicted by PRODIGAL (38) with the use of metaWRAP (32), and the *pI* of each ORF was calculate with ProPAS (39). The average *pI* of the entire gene pool as well as individual taxa were calculated from the average *pI* of proteins encoded on contigs of relevant (KRAKEN) taxonomy.

### Taxonomic turnover index (*TTI*)

The turnover indexes (*TTIs*) of each gene function (KO ID) represent the changes in relative abundances of the organismal strains (contigs) carrying them. For this purpose, the abundance of any given gene is assumed to be equal to the average abundance (coverage) of the contig that carries it. To calculate the *TTIs*, all contigs carrying genes of a given KEGG KO were identified, and the change in their relative abundances was calculated between two time-points of interest. Contig abundances from individual replicates were added up for each time point, then the *TTI* for each KEGG KO identifier was calculated from the weighted average of the absolute values of these changes (Equation 1). Importantly, this index does not measure the net change in abundance of each function, but instead quantifies the turnover in the organisms that carry it. Indeed, it is possible for the total abundance of a gene function to be carried by a completely new set of organisms, yet remain unchanged in total abundance. The *RIs* from all the KEGG functions were plotted and the difference in their distributions between the time points was computed with the Kolmogorov-Smirnov 2-sample test.

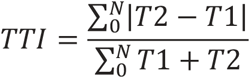

**Equation 1:** Formula calculating one function’s taxonomic turnover index *TTI*, where *T1* and *T2* are standardized abundances of a contig carrying that function in two samples, and *N* is the number of contigs carrying that functions.

### Shotgun statistical analysis

The significance in abundance changes of gene functions (i.e. KEGG KO identifiers), functional pathways (i.e. KEGG BRITE identifiers), and average *pI* of gene pools were estimated with a two-sided Student’s two-sample t-test. The relative similarity between groups of replicates (ordered by harvest dates) in terms of total pathway abundances and co-assembly contig abundances were computed by comparing Pearson correlations between samples. A Pearson correlation coefficient distance matrix was computed from all replicates, and a two-sided Student’s two-sample t-test was performed to evaluate the significance of the difference between the correlation distances. Differentially abundant KEGG (level 2) pathways were selected with a one-way ANOVA test (*p*<0.01, FDR<1%), and hierarchically clustered with Seaborn v0.8 (31) (method=‘average’, metric=‘euclidean’). The significance of the differences in distributions of *RIs* between pairs of time points, as well as differences in *pI* distributions of gene pool proteins were calculated with the Kolmogorov-Smirnov 2-sample test. Significance of MAG abundance, contig abundance, and pathway abundance clustering was determined with SigClust (nsim=1000, icovest=3) (40). Due to time considerations, the contig clustering test was limited to contigs over 5kbp in length, which were then subsampled randomly to 5000 contigs prior to the test.

## RESULTS

### High-order taxonomic composition and functional potential were temporarily perturbed after the rain

The halite communities were found to be sensitive to the acute perturbation from the rain at the end of 2015 (Fig. S2), as it induced a change in their taxonomic structure (Fig. 1). Practical considerations limited this longitudinal study to 4 samples collected of a 4-year period (2014-2017), with 2 time points before and 2 time points after the rain event. A second site was sampled 4 times after the rain, over 1 year. The average climate temperature during pre-rain sample collection was notably cooler (11°C-18°C) than that of 2016 and 2017 (17°C-25°C), which could have contributed to the shift described below. However, the recovery of the community composition in the following year despite higher temperatures suggests that the shift and recovery were primarily driven by the two rain events at the end of 2015.

**Fig. 1.**
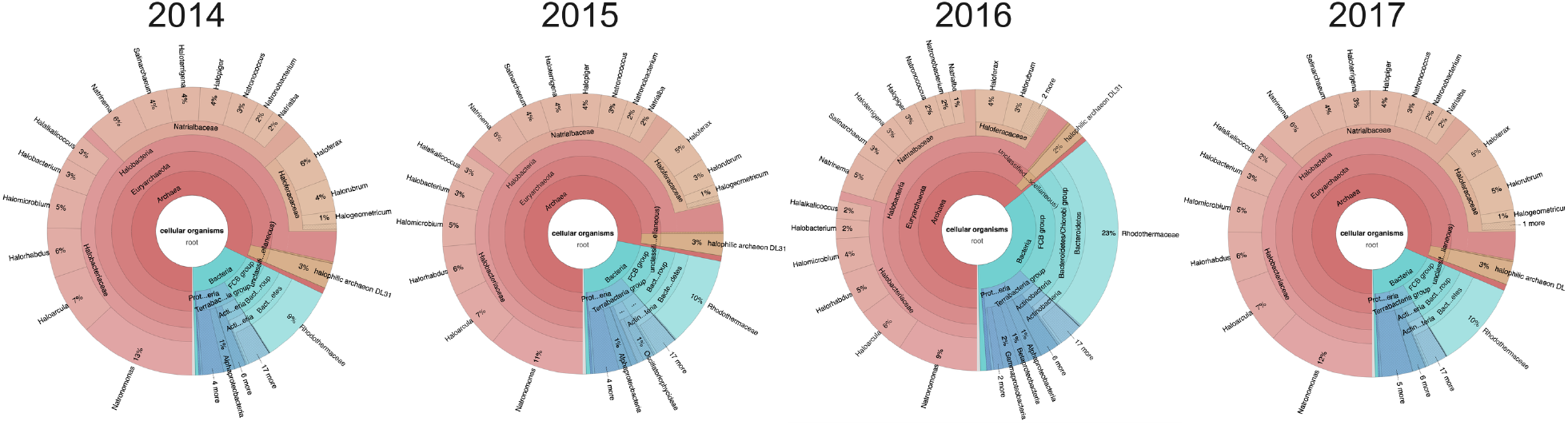
Average taxonomic composition of halite microbial communities from Site 1 before (2014, 2015) and after (2016, 2017) the rain event, estimated from whole metagenome reads with KRAKEN and visualized with KronaTools.

Weighted Unifrac analysis of the amplicon data, which compares the dissimilarity of communities based on weighted taxonomic composition, revealed that the halite communities were significantly different between time-points (PERMANOVA: *p*<0.001), with the taxonomic composition shifting following the rain. While the composition of the post-recovery (2017) communities was still significantly different from the pre-rain (2014 and 2015) samples (PERMANOVA: *p<*0.001), we found that they were more similar to each other than to the post-rain (2016) communities, suggesting a partial recovery in composition (two-sided t-tests of pairwise comparisons: *p*<0.0001; Fig. 2A, S3E). To investigate broad high-level taxonomic changes, we interrogated the community composition at the domain and phylum levels. At the domain level, the halite community structure shifted from an *Archaea*-dominated community before the rain (2014 and 2015) to a more balanced *Archaea-Bacteria* community 3-months after the rain (2016) (Fig. 1). The relative abundance of *Archaea* dropped significantly (two-sided t-tests: *p*<0.0001) in both 16S rRNA gene (Fig. 2B) and WMG sequencing. At the phylum level, we tracked changes in four taxa that constituted the majority of the community - Cyanobacteria, Bacteroidetes, Euryarchaeota (only represented by Halobacteria), and Chlorophyta (Data S1). While chloroplast 16S rRNA gene abundance is not necessarily indicative of the absolute abundance of algae, we know that there is only one alga in the halite community and that it contains a unique chloroplast (17), validating our use of chloroplast sequences as a proxy for relative algal abundances. All four taxa significantly shifted in abundance after the rain: Cyanobacteria, Chlorophyta and Bacteroidetes significantly increased in relative abundance following the rain, while the abundance of Halobacteria significantly decreased (Fig. 1, S3A-D; Data S1; two-sided t-tests: *p*<0.01). The abundances of these taxonomic groups partially recovered in the final sampling time-point (Fig. S3). To strengthen these observations of community changes, we conducted additional sampling after the rain with a higher temporal resolution at an alternate location (Site 2; Fig. S4, S5; Data S1). From 16S rRNA gene sequencing of this additional data set, we discovered gradual changes of domain (Fig. S4A) and some of the major phyla (Fig. S5) during the year after the rain, revealing the slow nature of this recovery process. Weighted Unifrac dissimilarity clustering of these samples (Fig. S4B) confirmed significant differences between the pre- (Feb 2016) and post-rain (Feb 2017) samples (PERMANOVA: *p<*0.001), however the intermediate time-points (Jul 2017 and Oct 2017) did not form distinct clusters and overlapped with the other samples.

**Fig. 2.**
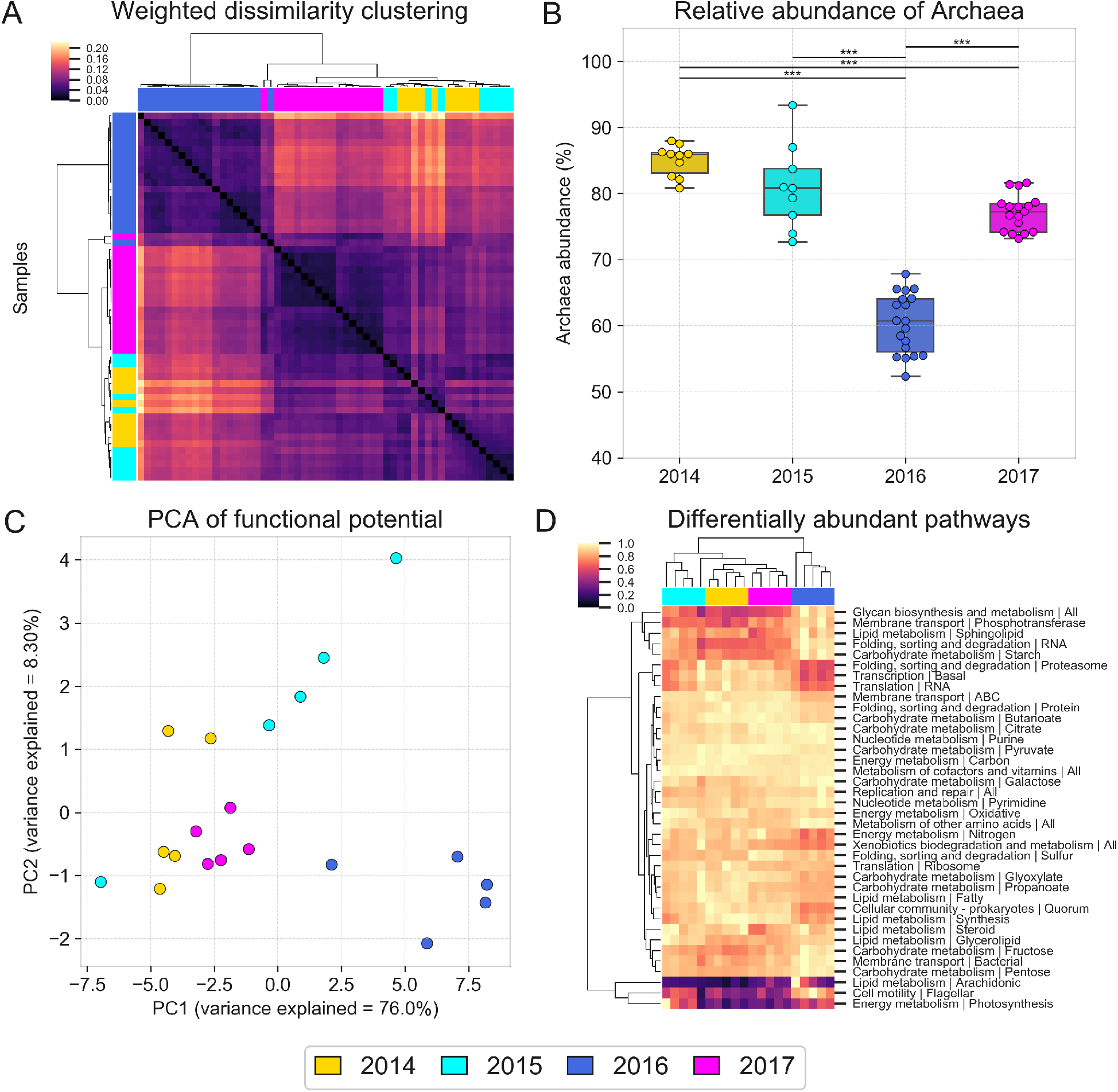
Halite microbial community taxonomic composition and functional potential over time. Taxonomic composition of halite microbiomes at each time point, shown by (A) hierarchical clustering (correlation metric) of the Weighted Unifrac dissimilarity matrix and (B) the average relative abundance of archaeal sequences, based on 16S rRNA gene amplicon sequencing. Error bars represent standard deviation; significance bars denote two tail t-test p-val<0.0001. The changes in functional potential of the halite communities is shown in (C) with a PCA of the abundance of KEGG pathways inferred from WMG co-assembly quantitation and (D) with hierarchical clustering (Euclidean metric) of differentially abundant pathways (ANOVA p<0.01, FDR=<1%), standardized to the maximum value in each row.

The functional potential of the community, determined by annotation of KEGG pathways in the WMG co-assembly, also significantly changed after the rain, although it is important to note that these estimates were only based on gene abundances. Consistent with the taxonomy-based clustering, samples from before the rain (2014 and 2015) were distinctly separate from samples collected shortly after the rain (2016; Fig. 2C). The KEGG pathway abundances in 2014 samples were better correlated with that of 2015 and 2017 samples than 2016 samples (two-sided t-tests of Pearson correlations: *p*<0.001). While the majority of functional pathways were present in similar abundances between replicates and time points, a number of pathways were differentially represented between time points (Fig. 2D; ANOVA test, *p*<0.01, FDR<1%). Of these, the majority were significantly over- or under-represented in the samples collected shortly after the rain (2016-02; SigClust 2-group significance: *p*<0.0001).

### Differences in salt adaptations likely drove changes in salt-in strategists

The most notable change in the functional composition of the community post-rain (2016) was an enrichment in proteins with a higher isoelectric point (*pI*), and a decrease in the potassium uptake potential (*trk* genes), both of which are hallmarks of salt-in strategists. We found that the *pI* of proteins encoded in community gene pool shifted significantly after the rain, favoring higher *pI* composition (Fig. 3A; KS 2-sample test: *p*<0.0001). Because of the significantly different *pI* distributions in the predicted proteins of Halobacteria (*pI*=5.04) and Bacteroidetes (*pI*=5.80; Fig. 3D; KS2-sample test: *p*<0.0001), the shift in their relative abundances resulted in the average *pI* of the community to significantly increase after the rain (two-sided t-test: *p*<0.01; Fig. 3B). Consistent with salt-in adaptations, we also found that the average potassium uptake potential (estimated from *trk* gene abundances) significantly decreased after the rain (Fig. 3C). Interestingly, both the shift in the average protein pool *pI* and the change in potassium uptake potential were also observed within the highly heterogeneous Halobacteria class (Fig. 3E, F).

**Fig. 3.**
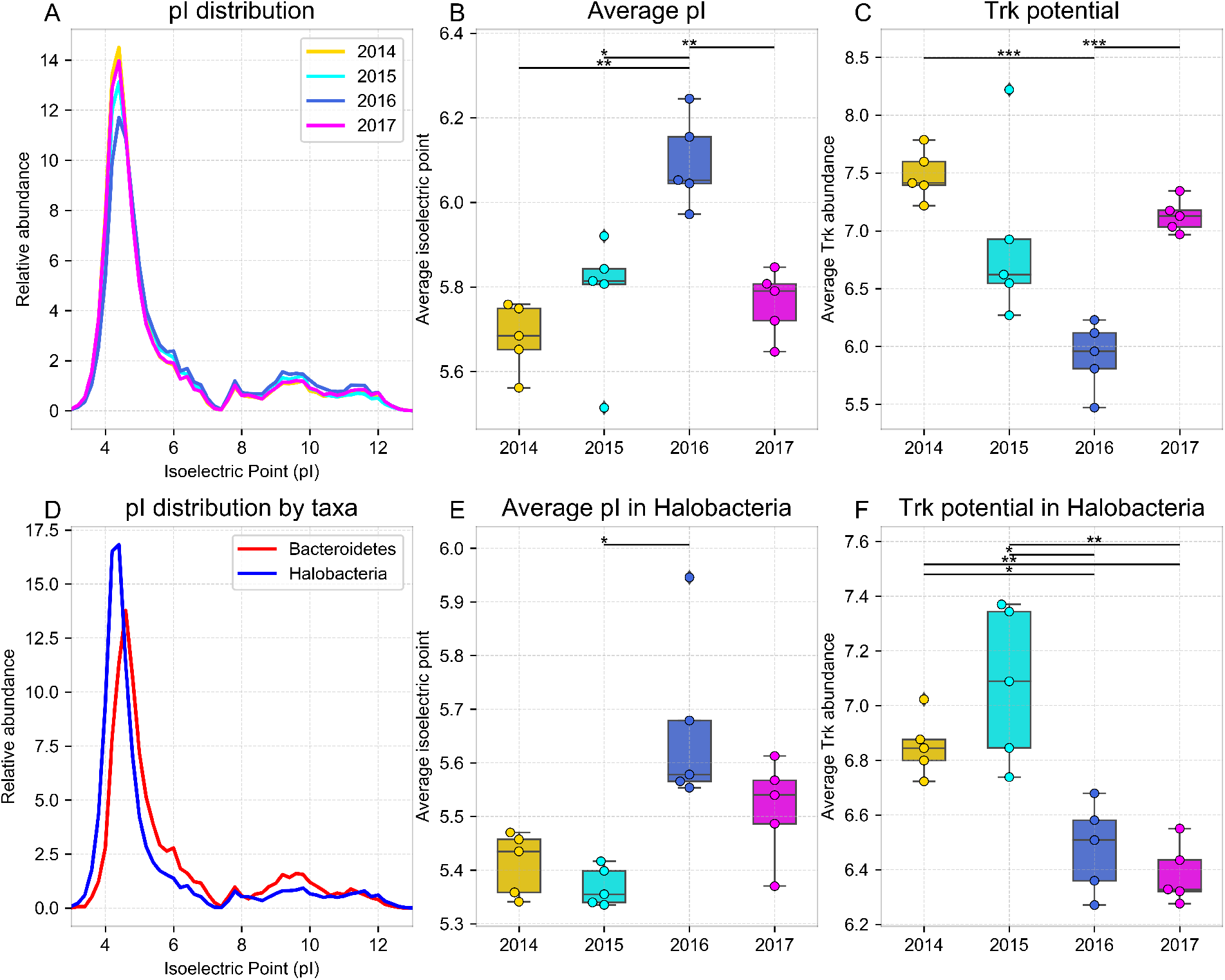
Differences in predicted protein isoelectric points and potassium uptake potential over time. Analysis of the isoelectric points (pI) of proteins encoded in replicates of WMG assemblies from samples harvested at different dates, showing (A) the overall weighted distribution of the protein pIs, and the weighted average pI of proteins encoded in (B) all contigs and (E) only Halobacteria contigs. (D) pI distribution of predicted proteins encoded in Bacteroidetes and Halobacteria contigs. Average potassium uptake potential across time point samples inferred from trk gene relative abundance and quantified in (C) all contigs and (F) only Halobacteria contigs. Error bars represent standard deviation; significance bars represent group significance based on a two tail t-test, and stars denote the p-value thresholds (*=0.01, **=0.001, ***=0.0001).

### Fine-scale taxonomic compositional shift after the rain

While changes in overall taxonomic composition (domain and phylum levels) of the halite communities were transient (Fig. 2A,B), we surprisingly found that their fine-scale composition (individual OTUs and contigs) did not recover. Samples collected at different dates were significantly different in terms of presence or absence of operational taxonomic units (97%OTUs), as measured by the Unweighted Unifrac dissimilarity index (PERMANOVA: *p*<0.001), with samples harvested shortly after the rain (2016) being more distant from pre-rain samples than they were from each other (two-sided t-test: *p*<0.0001). We found that the community did not return to its initial state after the perturbation, as the post-recovery samples (2017) clustered together with post-rain (2016) samples (Fig. 4A), and were less distant to 2016 samples than to the pre-rain samples (two-sided t-test: *p*<0.0001). The altered OTU composition of the community, shown with Unweighted Unifrac clustering, contrasts with the successful recovery of the higher-order taxonomic structure, as shown with Weighted Unifrac dissimilarity clustering (Fig. 2A).

**Fig. 4.**
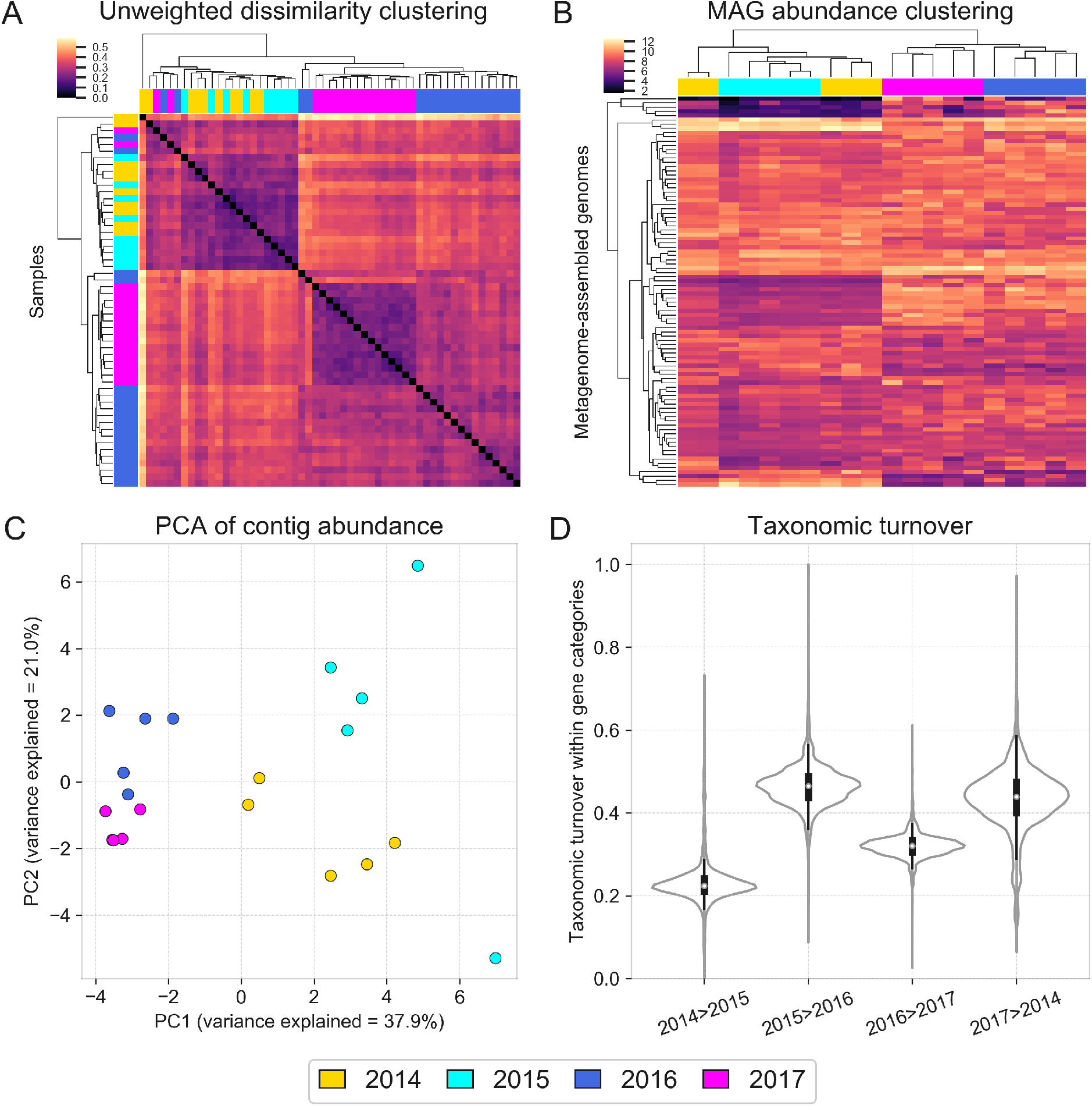
Fine-scale taxonomic composition shifts across time. Fine-scale compositional changes of halite communities over time shown with (A) hierarchical clustering (correlation metric) of an Unweighted Unifrac dissimilarity matrix (based on 16S rRNA gene amplicon sequencing), (B) hierarchical clustering (Euclidean metric) of standardized MAG abundances, (C) PCA of co-assembly contig abundances, and (D) weighted distributions of taxonomic turnover (TTI) of functional niches between time points. The TTI of each functional category estimates the changes in organisms that encode it (see Methods).

The shift in the community’s fine-scale membership was validated with WMG sequencing at the scale of individual contig abundances (Fig. S6). Based on contig read coverage across samples, we found that all post-rain samples clustered away from pre-rain samples (Fig. 4C; SigClust 2-group significance: *p*<0.01). Additionally, pairwise Pearson correlation comparison confirmed that contig abundances of post-rain samples were better correlated with each other than with that of pre-rain samples (two-sided t-test: *p*<0.0001). These contig-level turnover dynamics were additionally investigated with individually recovered metagenome-assembled genomes (MAGs). 91 high-quality MAGs (>70% completion, <5% contamination; Data S2) were reconstructed with metaWRAP (32) and their abundances were tracked between samples. Pearson correlation comparison (two-sided t-test: *p*<0.0001) and group significance analysis (SigClust 2-group significance: *p*<0.01) confirmed the permanent shift of the fine-scale taxa composition after the rain (Fig. 4B). While the fine-scale composition of the community did change during the post-rain recovery between 2016 and 2017, the resulting shift was more moderate when compared to the more drastic taxonomic shift immediately following the rain. This contrasts with the near-complete recovery of the overall functional potential of the community (Fig. 2C,D). Additionally, two conditionally rare taxa (41) of *Cyanobacteria* that were previously reported in only a small fraction of halite nodules (18), were found in high abundances in most of the samples after the rain (Fig. S7). Surprisingly, we found no correlation between the functional potentials of the MAGs and their survival after the rain, suggesting that this shift was a stochastic process. These results indicate that while the abundances of higher-order taxonomic ranks recovered to the pre-rain state, the fine-grain taxonomy of the community has been permanently reshuffled.

### The rain disrupted taxonomic membership of potential functional niches

To investigate the basis of the functional potential shift of the halite community after the rain, we introduced a taxonomic turnover index (*TTI*), which quantifies the turnover of strains (estimated from contigs) contributing to each community function. To compute the *TTI*, genes from each KEGG Orthology identifier were catalogued and their abundances in each sample estimated from the read coverage of the contig that they were on. The absolute value average of the change in contig abundances that carry a given function between two samples represents the degree of taxonomic turnover within that functional category (see Methods). A relatively high *TTI* for a given community function indicates that it is carried by different community members between two samples, but does not necessarily imply a high net change in its total abundance in the samples. Therefore, the distribution in *TTIs* for all functions between two time-points quantifies changes in niche representation over that time (Fig. 4D). However, because these results are based solely on functional potential prediction from gene abundances, it should be noted that our estimations of the functional landscape could be significantly altered by compensatory transcriptional and translational processed, and functional rates. The turnover following the rain (2015 to 2016) was significantly higher than the baseline taxonomic shift prior to the rain (2014 to 2015; KS 2-sample test: *p*<0.0001), indicating that the same functional pathways were being carried on a different set of contigs. However, the shift in functional niche membership during the recovery phase (2016 to 2017) was low compared to the post-rain shift, indicating that the taxonomic membership did not return to its initial state. These findings indicate that functional redundancy of community members ensured a robust functional landscape in the halite microbial communities despite changes in the fine-scale taxonomic membership. Interestingly, this shift did not notably affect the overall functional diversity of the samples, as seen from lack of a significant difference between the total number of unique gene functions found in each time-point (two-sided t-tests: *p*>0.05).

## DISCUSSION

The response and recovery of the halite microbiome, a sensitive extremophile ecosystem, provided the opportunity to characterize the response dynamics of a natural community to changing environmental conditions. While low sampling frequency limits the temporal resolution of this study, our evidence suggests that the 2015 rainfall required major adaptations in the extreme halophiles found within the halite nodules of Salar Grande. The shift in the observed taxonomic composition following the rain was noteworthy not only in the context of this study but also when comparing with previous studies of this area in 2013 (17). The surviving community was comprised of organisms with higher average isoelectric points (*pI*) of their predicted proteins and lower potassium uptake potential. This was significant because high potassium uptake is a strategy used by salt-in strategists to balance high external salt concentrations, while the low *pI* of their proteome allows them to function in the high-potassium intracellular environment (21, 42). Our reported average isoelectric points for the two dominant salt-in strategists in this system – 5.80 (Bacteroidetes) and 5.04 (Halobacteria) – were similar to those previously documented for these taxa – 5.92 and 5.03, respectively (19). It is also well documented that acid-shifted proteomes is also an adaptation in salt-in strategists to increasing salt in the environments (43, 44). The changes in *pI* and potassium uptake potential we observed after the rain suggest that the rain temporarily decreased the salt concentrations within the colonized pores (23, 45), rapidly changing the osmotic conditions within. We hypothesize that this led to a mass death event of organisms poorly adapted to large osmotic changes immediately following the rain, while giving others an advantage.

The taxonomic shifts at the contig level were likely driven by neutral (i.e. random) processes (46, 47) resulting from the halite re-colonization, rather than deterministic processes associated with adaptation to the rain. These stochastic dynamics, similar to those governing the initial colonization of halite nodules, resulted in high inter-nodule taxonomic diversity (18) while the functional states remained. We suggest that each nodule was stochastically colonized by random draw, from the seed bank, of competitively equivalent organisms. A seed bank is a diverse genetic reservoir consisting of a large collection of low-abundance organisms (1, 48) that might be critical for microbiome functioning, particularly following prolonged unchanging environmental conditions such as the past 13 years prior to the rain in northern Atacama. Seed banks conserve genetic and functional diversity, which in turn allows for rapid adaptation and restructuring of the microbial community following a drastic perturbation.

While our methods cannot differentiate the DNA of living organisms from relic DNA present in the halite nodule (49), it is unlikely that the observed compositional shift after the rain was an artifact of relic DNA turnover. Indeed, it is improbable that the 24.2mm of rain was sufficient to wash away relic DNA from within the nodules. Similarly, the rain itself was probably not a major contributor to the sequenced DNA since we did not detect non-halophilic organisms that are likely to be found in atmospheric microbiomes (50).

The halite microbiome was able to recover from this catastrophic event, however, the effects of the perturbation lasted remarkably long (months), in contrast with studies in other desert systems where much quicker recoveries were documented (weeks) (12). The higher temporal resolution in the time series at additional sampling Site 2 especially highlights the slow-growing nature of these extremophiles and suggests that the immediate effects of the rain on the halite community may have been even more dramatic than what we observed 3-months post-rain (17, 51). Fifteen months post-rain, the community was comprised of an entirely new set of organisms but its functional potential recovered to a pre-rain state, suggesting that the community taxonomic structure entered an alternative equilibrium state during the recovery period (4, 11). The functional consistency of a community, disconnected from taxonomic variance, has previously been documented in a variety of microbiomes and stems from functional redundancy of closely related taxa (6, 7, 52, 53). In particular, isolated microbiomes such as miniature aquatic ecosystems found in bromeliad rosettes (similarly isolated as the halite nodules) appear to converge on identical functional landscapes through mechanisms such as stoichiometric balancing between metabolic pathways, despite great inter-community taxonomic diversity (54, 55).

The pre-rain (2014) and recovered (2017) communities were very similar in terms of their functionally potential, while the intermediate state (2016) was very distinct (Fig. 2C, D). Therefore, the two compositional shifts that the halite microbiomes underwent following the rain – the initial response (2015-2016) and subsequent recovery (2016-2017) – resulted in a similar magnitude of change to the overall functional potential of the community. Taxonomically however, the two shifts were fundamentally distinct, as the individual taxa membership was drastically changed during the initial response to the rain but stayed unchanged during the recovery (Fig. 4B, C).

The two different mechanisms by which the halite communities achieved almost identical net change in their functional potential as they entered and then exited their intermediate state (11, 12) offered a uniquely detailed view of microbial adaptation dynamics. These two types responses, or modes, allowed for inference of a general microbiome adaptation model, which can be potentially applied to explain and predict the taxonomic and functional flux in other ecosystems following major environmental changes (Fig. 5). The first mode (*Type I*; Fig. 5A) is a community shift, resulting from adaptations to an acute major perturbation. In the halite nodules, the rain presented a major stress on the pre-existing communities by temporarily lowering external osmotic conditions and exerting a strong selective pressure on the salt-in strategists. This produced gaps in existing functional niches and presented an opportunity for new organisms from the seed bank to come in through niche intrusion (56). The *Type I* shift is driven by neutral (random) processes characterized by changes in fine-scale (i.e. strains) taxonomic composition, which results in a high taxonomic turnover index (*TTI*=0.89±0.12 in the model).

**Fig. 5.**
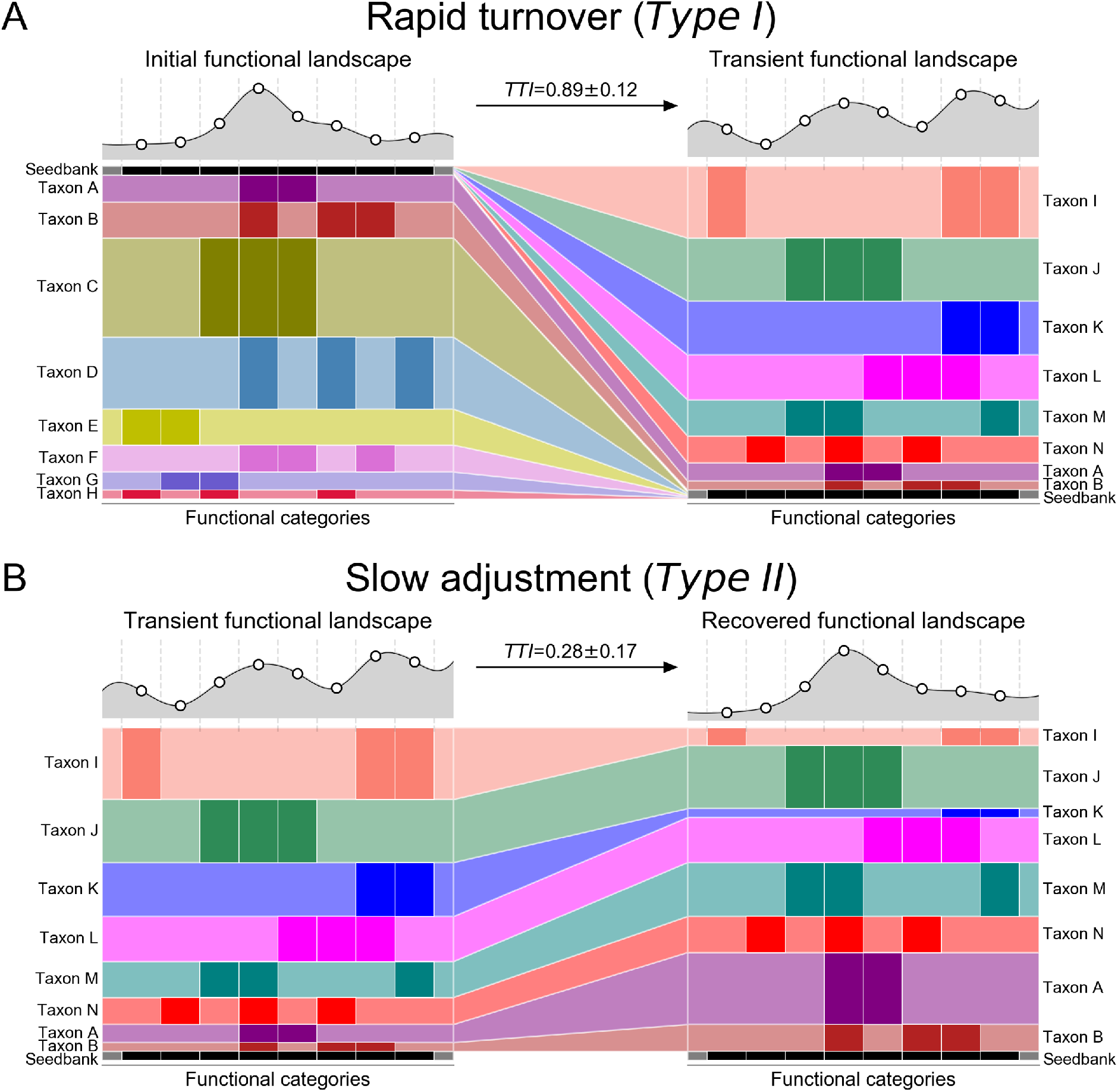
Microbial community resilience model. Models of a microbiome adapting its functional potential in response to changing environmental conditions either with (A) a rapid shift of the community’s taxonomic composition resulting from new organisms from the seed bank displacing previously dominant taxa through niche intrusion (as seen in the initial shock from the rainfall), or with (B) a gradual adjustment in relative abundance of major taxa (as seen in the halite community recovery). On the y-axis, the vertical spread represents the abundance of a given taxon (A through H) and on the x-axis, darker colored bars show which functional category is encoded in their genomes. The seedbank (black bars) represents rare taxa in the community. The functional landscape curves at the top of each figure visualize the relative total abundance of each functional category, calculated by adding the abundances of the organisms that carry that function. Taxonomic turnover (TTI) rates were calculated for each model community in (A) and (B).

The second mode (*Type II*; Fig. 5B) is an adjustment in existing community structure, and results from gradual changes in environmental conditions. After the rain passed and the osmotic conditions within the halite nodules returned to their initial levels, the halite community gradually returned to its previous functional potential. However, because there were no major stress events to reset the strain composition of the communities, the newly dominant strains remained relatively unchanged during the recovery period. Instead, the functional potential of the community is achieved through gradual changes in relative abundances of major taxa (Fig. 2, S4, S5), the strain composition of which remained unchanged. The taxonomic mechanism behind the *Type II* response is relatively deterministic, as the relative abundances of currently dominant taxa is adjusted based on fitness under the new selective pressures, preventing new organisms to take over. As a result, the strain composition of these major taxa remain largely unchanged, resulting in a low taxonomic turnover index (*TTI*=0.28±0.17 in the model). In the halite microbiome, the *Type I* and a *Type II* shifts occurred in succession, leading the community first through an unstable intermediate state and then into an alternate equilibrium state (4). This intermediate dis-equilibrium intermediate has been reported in a number of communities after disaster events (57) or antibiotic administration (56, 58), but until now was difficult to investigate closely in natural ecosystems because of compounding complexity and fast microbial growth rates (1, 4). We postulate that *Type I* and *Type II* shifts observed in our model microbiome are integral to analogous structural rearrangement in other systems.

It is important to note that *Type I* and *Type II* functional shifts do not necessarily follow one another. If the initial environmental conditions are not re-established after a perturbation, such as after a permanent introduction of irrigation to desiccated soils, a *Type I* shift will most likely be the main mechanism for community adaptation, driven by the changes in environmental conditions. Alternatively, in systems where environmental conditions shift gradually, such as aquatic microbiomes during seasonal changes, Type II shifts will likely drive the changes in the community’s functional potential. We propose that *TTI* measurements of such shifts may be useful in future studies to categorize such dynamics.

In conclusion, the tractable nature of our model microbiome allowed us to extrapolate general mechanisms of community response and resilience to acute shock. We demonstrated that a major disturbance can result in stochastic re-population of the community’s functional niches, forcing a microbial community structure into an unstable intermediate. During the succeeding recovery period, the newly dominant taxa adjust in abundance to reproduce the initial functional potential, allowing the community to enter an alternative equilibrium. Understanding the mechanisms behind the response and recovery components of microbial perturbation responses are vital to generally model and predict the taxonomic and functional flux of ecosystems following natural and man-made ecological disasters. Our proposed characterization and quantitation of two types of community shifts and our two-step model for community resilience can provide a framework for future work in predictive modeling of microbial communities.

## General

We thank Mike Sauria and Boris Brenerman for data analysis suggestions, Diego Gelsinger and Alexandra Galetovic for help with fieldwork, and Sarah Preheim, Michael Schatz and Vadim Uritsky for help in manuscript editing.

## Author contributions

GU, JT, and JD conceptualized and designed the study; GU, JD, BGS and AD collected in-field samples; BG organized and funded field expeditions; SG and AM processed and sequenced samples; GU analyzed the data and wrote the manuscript; JT and JD edited the manuscript.

## Data and materials availability

Raw sequencing data is available from the National Centre for Biotechnology Information under project ID PRJNA484015. All analysis pipelines, processed data, analysis and visualization scripts, and reconstructed MAGs are available at https://github.com/ursky/timeline_paper. The metagenome co-assembly and functional annotation are available from the JGI Genome Portal under IMG taxon OID 3300027982.

**Supplementary information is available online.**

## REFERENCES

1. Shade A, Peter H, Allison SD, Baho DL, Berga M, Burgmann H, et al. Fundamentals of microbial community resistance and resilience. Front Microbiol. 2012;3:417.

2. Raymond F, Deraspe M, Boissinot M, Bergeron MG, Corbeil J. Partial recovery of microbiomes after antibiotic treatment. Gut Microbes. 2016;7(5):428–34.

3. David LA, Maurice CF, Carmody RN, Gootenberg DB, Button JE, Wolfe BE, et al. Diet rapidly and reproducibly alters the human gut microbiome. Nature. 2014;505(7484):559–63.

4. Scheffer M, Carpenter S, Foley JA, Folke C, Walker B. Catastrophic shifts in ecosystems. Nature. 2001;413(6856):591–6.

5. Jurburg SD, Nunes I, Brejnrod A, Jacquiod S, Prieme A, Sorensen SJ, et al. Legacy Effects on the Recovery of Soil Bacterial Communities from Extreme Temperature Perturbation. Front Microbiol. 2017;8:1832.

6. Lozupone CA, Stombaugh JI, Gordon JI, Jansson JK, Knight R. Diversity, stability and resilience of the human gut microbiota. Nature. 2012;489(7415):220–30.

7. Goldford JE, Lu N, Bajic D, Estrela S, Tikhonov M, Sanchez-Gorostiaga A, et al. Emergent Simplicity in Microbial Community Assembly. bioRxiv. 2017.

8. Palleja A, Mikkelsen KH, Forslund SK, Kashani A, Allin KH, Nielsen T, et al. Recovery of gut microbiota of healthy adults following antibiotic exposure. Nat Microbiol. 2018;3(11):1255–65.

9. Thiemann S, Smit N, Strowig T. Antibiotics and the Intestinal Microbiome : Individual Responses, Resilience of the Ecosystem, and the Susceptibility to Infections. Curr Top Microbiol Immunol. 2016;398:123–46.

10. Jernberg C, Lofmark S, Edlund C, Jansson JK. Long-term impacts of antibiotic exposure on the human intestinal microbiota. Microbiology. 2010;156(Pt 11):3216–23.

11. Allison SD, Martiny JB. Colloquium paper: resistance, resilience, and redundancy in microbial communities. Proc Natl Acad Sci U S A. 2008;105 Suppl 1:11512–9.

12. Armstrong A, Valverde A, Ramond JB, Makhalanyane TP, Jansson JK, Hopkins DW, et al. Temporal dynamics of hot desert microbial communities reveal structural and functional responses to water input. Sci Rep. 2016;6:34434.

13. McKay CP, Friedmann EI, Gomez-Silva B, Caceres-Villanueva L, Andersen DT, Landheim R. Temperature and moisture conditions for life in the extreme arid region of the Atacama desert: four years of observations including the El Niño of 1997-1998. Astrobiology. 2003;3(2):393–406.

14. Bozkurt D, Rondanelli R, Garreaud R, Arriagada A. Impact of Warmer Eastern Tropical Pacific SST on the March 2015 Atacama Floods. Monthly Weather Review. 2016;144(11):4441–60.

15. Wierzchos J, Casero MC, Artieda O, Ascaso C. Endolithic microbial habitats as refuges for life in polyextreme environment of the Atacama Desert. Current Opinion in Microbiology. 2018;43:124–31.

16. Robinson CK, Wierzchos J, Black C, Crits-Christoph A, Ma B, Ravel J, et al. Microbial diversity and the presence of algae in halite endolithic communities are correlated to atmospheric moisture in the hyper-arid zone of the Atacama Desert. Environ Microbiol. 2015;17:299–315.

17. Crits-Christoph A, Gelsinger DR, Ma B, Wierzchos J, Ravel J, Davila A, et al. Functional interactions of archaea, bacteria and viruses in a hypersaline endolithic community. Environ Microbiol. 2016;18(6):2064–77.

18. Finstad KM, Probst AJ, Thomas BC, Andersen GL, Demergasso C, Echeverria A, et al. Microbial Community Structure and the Persistence of Cyanobacterial Populations in Salt Crusts of the Hyperarid Atacama Desert from Genome-Resolved Metagenomics. Front Microbiol. 2017;8:1435.

19. Mongodin EF, Nelson KE, Daugherty S, DeBoy RT, Wister J, Khouri H, et al. The genome of Salinibacter ruber: Convergence and gene exchange among hyperhalophilic bacteria and archaea. PNAS. 2005:0509073102.

20. Monard C, Gantner S, Bertilsson S, Hallin S, Stenlid J. Habitat generalists and specialists in microbial communities across a terrestrial-freshwater gradient. Sci Rep. 2016;6:37719.

21. Oren A. Life at high salt concentrations, intracellular KCl concentrations, and acidic proteomes. Front Microbiol. 2013;4:315.

22. Thombre RS, Shinde VD, Oke RS, Dhar SK, Shouche YS. Biology and survival of extremely halophilic archaeon Haloarcula marismortui RR12 isolated from Mumbai salterns, India in response to salinity stress. Sci Rep. 2016;6:25642.

23. Davila AF, Hawes I, Araya JG, Gelsinger DR, DiRuggiero J, Ascaso C, et al. In situ metabolism in halite endolithic microbial communities of the hyperarid Atacama Desert. Front Microbiol. 2015;6:1035.

24. Weather Underground 2019 [03/19/19]. Available from: https://www.wunderground.com/history/monthly/cl/iquique/SCDA.

25. Schulz N, Boisier JP, Aceituno P. Climate change along the arid coast of northern Chile. International Journal of Climatology. 2012;32(12):1803–14.

26. Azua-Bustos A, Fairen AG, Gonzalez-Silva C, Ascaso C, Carrizo D, Fernandez-Martinez MA, et al. Unprecedented rains decimate surface microbial communities in the hyperarid core of the Atacama Desert. Sci Rep. 2018;8(1):16706.

27. Needham DM, Fuhrman JA. Pronounced daily succession of phytoplankton, archaea and bacteria following a spring bloom. Nat Microbiol. 2016;1:16005.

28. Caporaso JG, Kuczynski J, Stombaugh J, Bittinger K, Bushman FD, Costello EK, et al. QIIME allows analysis of high-throughput community sequencing data. Nat Methods. 2010;7(5):335–6.

29. Quast C, Pruesse E, Yilmaz P, Gerken J, Schweer T, Yarza P, et al. The SILVA ribosomal RNA gene database project: improved data processing and web-based tools. Nucleic Acids Res. 2013;41(Database issue):D590–6.

30. Edgar RC. Search and clustering orders of magnitude faster than BLAST. Bioinformatics. 2010;26:2460–1.

31. Waskom M, Botvinnik O, O’Kane D, Hobson P, Lukauskas S, Gemperline DC, et al. Seaborn. 0.8.1 ed: GitHub; 2017. p. https://github.com/mwaskom/seaborn.

32. Uritskiy GV, DiRuggiero J, Taylor J. MetaWRAP-a flexible pipeline for genome-resolved metagenomic data analysis. Microbiome. 2018;6(1):158.

33. Wood DE, Salzberg SL. Kraken: ultrafast metagenomic sequence classification using exact alignments. Genome Biol. 2014;15(3):R46.

34. Nurk S, Meleshko D, Korobeynikov A, Pevzner PA. metaSPAdes: a new versatile metagenomic assembler. Genome Res. 2017;27(5):824–34.

35. Patro R, Duggal G, Love MI, Irizarry RA, Kingsford C. Salmon provides fast and bias-aware quantification of transcript expression. Nat Methods. 2017;14(4):417–9.

36. Chen IA, Markowitz VM, Chu K, Palaniappan K, Szeto E, Pillay M, et al. IMG/M: integrated genome and metagenome comparative data analysis system. Nucleic Acids Res. 2017;45(D1):D507–D16.

37. Kanehisa M, Sato Y, Kawashima M, Furumichi M, Tanabe M. KEGG as a reference resource for gene and protein annotation. Nucleic Acids Res. 2016;44(D1):D457–62.

38. Hyatt D, Chen GL, Locascio PF, Land ML, Larimer FW, Hauser LJ. Prodigal: prokaryotic gene recognition and translation initiation site identification. BMC Bioinformatics. 2010;11:119.

39. Wu S, Zhu Y. ProPAS: standalone software to analyze protein properties. Bioinformation. 2012;8(3):167–9.

40. Liu Y, Hayes DN, Nobel A, Marron JS. Statistical Significance of Clustering for High-Dimension, Low-Sample Size Data. Journal of the American Statistical Association. 2008;103(483):1281–93.

41. Shade A, Jones SE, Caporaso JG, Handelsman J, Knight R, Fierer N, et al. Conditionally Rare Taxa Disproportionately Contribute to Temporal Changes in Microbial Diversity. Mbio. 2014;5(4).

42. Paul S, Bag SK, Das S, Harvill ET, Dutta C. Molecular signature of hypersaline adaptation: insights from genome and proteome composition of halophilic prokaryotes. Genome Biol. 2008;9(4):R70.

43. Kiraga J, Mackiewicz P, Mackiewicz D, Kowalczuk M, Biecek P, Polak N, et al. The relationships between the isoelectric point and: length of proteins, taxonomy and ecology of organisms. BMC Genomics. 2007;8:163.

44. Elevi Bardavid R, Oren A. Acid-shifted isoelectric point profiles of the proteins in a hypersaline microbial mat: an adaptation to life at high salt concentrations? Extremophiles. 2012;16(5):787–92.

45. Finstad K, Pfeiffer M, McNicol G, Barnes J, Demergasso C, Chong G, et al. Rates and geochemical processes of soil and salt crust formation in Salars of the Atacama Desert, Chile. Geoderma. 2016;284:57–72.

46. Hubbell SP. The Unified Neutral Theory of Biodiversity and Biogeography.. Princeton: New Jersey: Princeton Univ Press; 2001.

47. Li L, Ma ZS. Testing the Neutral Theory of Biodiversity with Human Microbiome Datasets. Sci Rep. 2016;6:31448.

48. Lennon JT, Jones SE. Microbial seed banks: the ecological and evolutionary implications of dormancy. Nat Rev Microbiol. 2011;9(2):119–30.

49. Schulze-Makuch D, Wagner D, Kounaves SP, Mangelsdorf K, Devine KG, de Vera JP, et al. Transitory microbial habitat in the hyperarid Atacama Desert. Proc Natl Acad Sci U S A. 2018;115(11):2670–5.

50. Caliz J, Triado-Margarit X, Camarero L, Casamayor EO. A long-term survey unveils strong seasonal patterns in the airborne microbiome coupled to general and regional atmospheric circulations. Proc Natl Acad Sci U S A. 2018;115(48):12229–34.

51. Ziolkowski LA, Wierzchos J, Davila AF, Slater GF. Radiocarbon evidence of active endolithic microbial communities in the hyper-arid core of the Atacama Desert,. Astrobiology. 2013;13:607–16.

52. Eng A, Borenstein E. Taxa-function robustness in microbial communities. Microbiome. 2018;6(1):45.

53. Nie Y, Zhao JY, Tang YQ, Guo P, Yang Y, Wu XL, et al. Species Divergence vs. Functional Convergence Characterizes Crude Oil Microbial Community Assembly. Front Microbiol. 2016;7:1254.

54. Louca S, Jacques SMS, Pires APF, Leal JS, Srivastava DS, Parfrey LW, et al. High taxonomic variability despite stable functional structure across microbial communities. Nat Ecol Evol. 2016;1(1):15.

55. Louca S, Polz MF, Mazel F, Albright MBN, Huber JA, O–Connor MI, et al. Function and functional redundancy in microbial systems. Nature Ecology & Evolution. 2018;2(6):936–43.

56. Modi SR, Collins JJ, Relman DA. Antibiotics and the gut microbiota. J Clin Invest. 2014;124(10):4212–8.

57. Rodriguez RL, Overholt WA, Hagan C, Huettel M, Kostka JE, Konstantinidis KT. Microbial community successional patterns in beach sands impacted by the Deepwater Horizon oil spill. ISME J. 2015;9(9):1928–40.

58. Sommer MO, Dantas G. Antibiotics and the resistant microbiome. Curr Opin Microbiol. 2011;14(5):556–63.

