## supplementary material for "Halophilic microbial community compositional shift after a rare rainfall in the Atacama Desert"

### Manuscript

**Supplementary figures:**

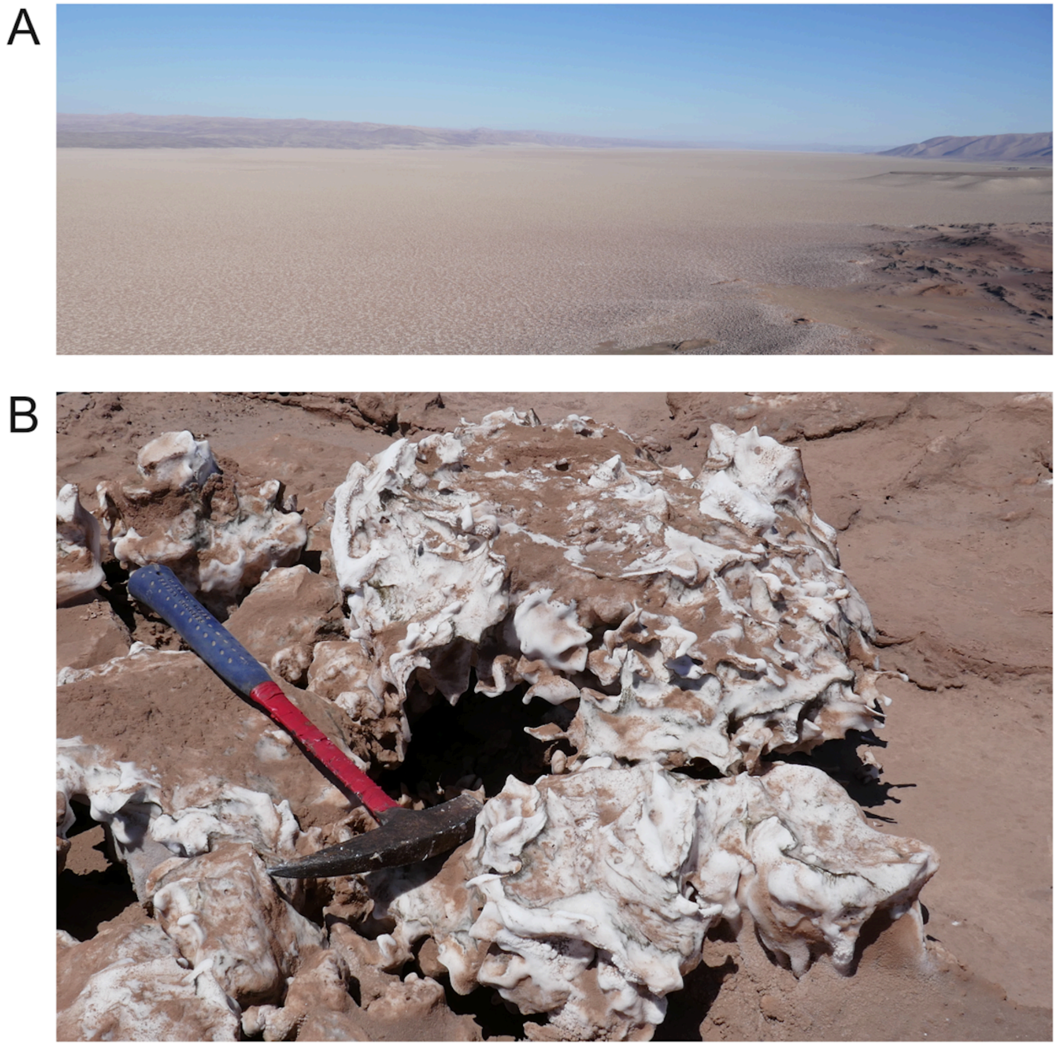

**Fig. S1.** Salar Grande landscape and halite nodules. (A) Aerial view of the evaporitic basin of Salar Grande, 5 km wide and 45 km long (N-S direction). (B) Halite nodules (salt rocks) 20 to 50 cm in size.

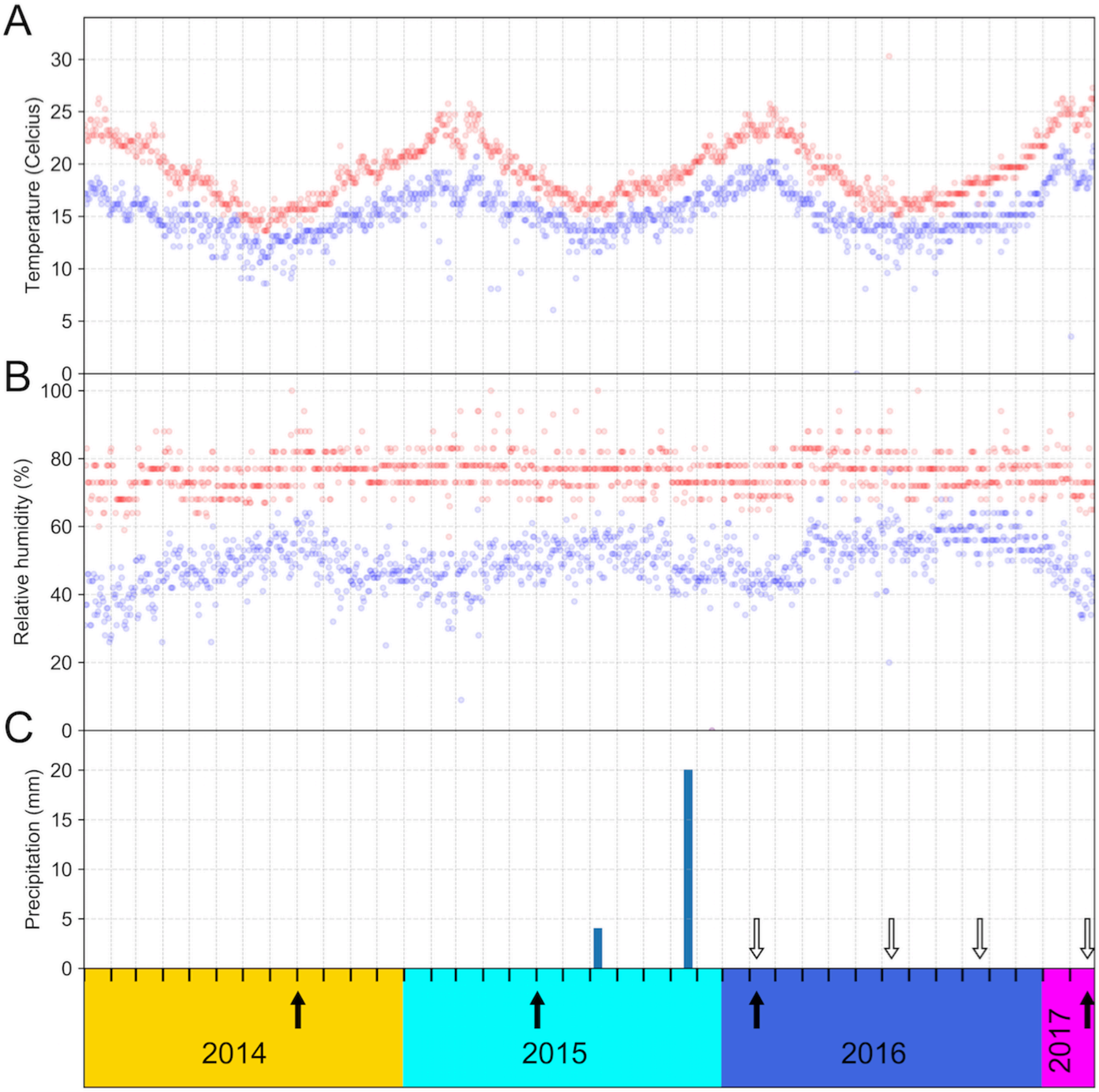


**Fig. S2.** Regional climate data from the Diego Aracena International Airport weather station, 40km North-West of Salar Grande. The maximum (red) and minimum (blue) temperature (A) and relative humidity (B) values, and total daily precipitation (C), are plotted for each date along the x-axis. Colors denote the year (2014-2017), x-ticks denote months, black arrows show the main sampling dates at Site 1, and white arrows show the sampling dates at Site 2.

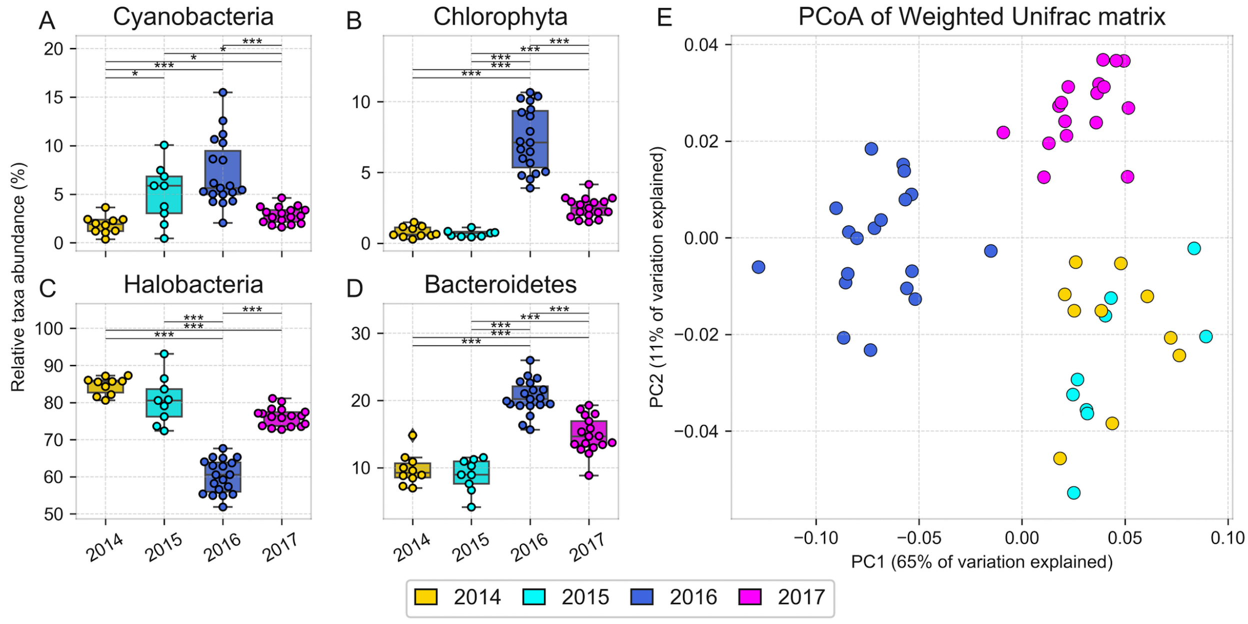


**Fig. S3.** Taxonomic composition of halite nodules from Site 1 over time inferred from 16S rRNA gene sequences clustered into OTUs at 97% identity and visualized through (A-D) relative abundance of the dominant phyla (Chloroplast was used as a proxy for Chlorophyta and Halobacteria was the only class of Euyarchaeota) whose abundance significantly shifted after the rain and a (E) PCoA plot of a Weighted Unifrac dissimilarity matrix comparing taxonomic composition. Error bars represent standard deviation; significance bars represent group significance based on a two tail t-test, and stars denote the p-value thresholds (*=0.01, **=0.001, ***=0.0001).

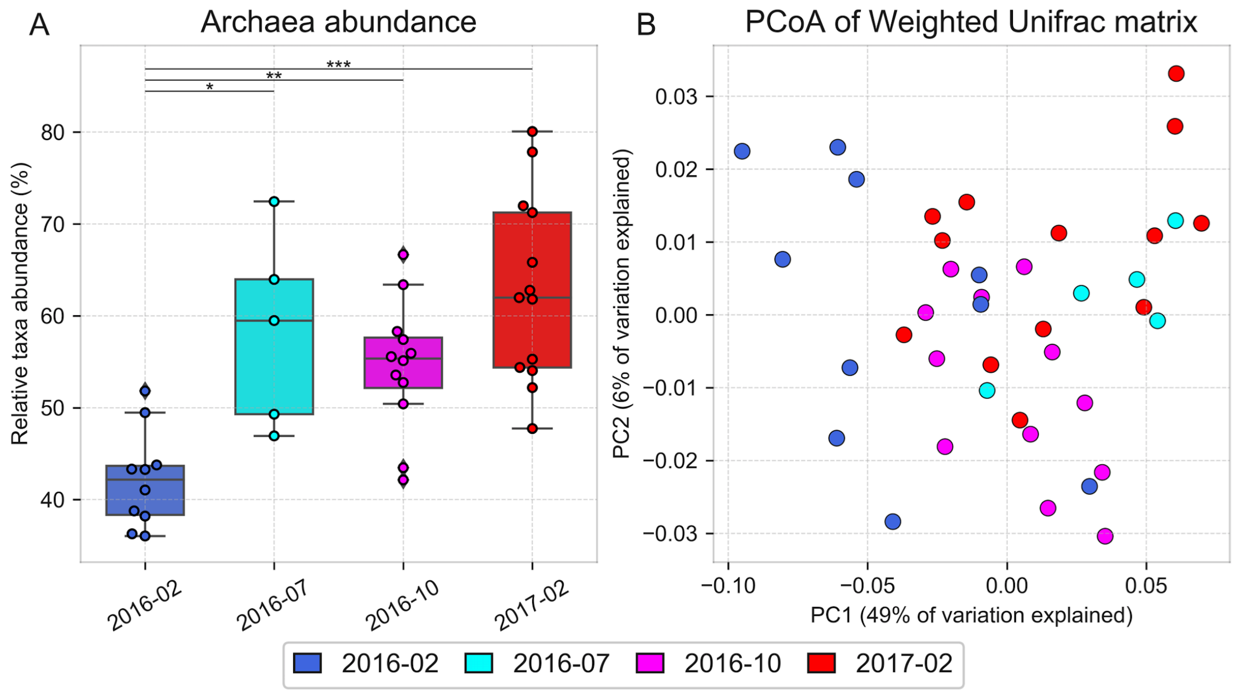


**Fig. S4.** Taxonomic composition of halite nodules harvested post-rain from Site 2 over time, inferred from 16S rRNA gene sequences clustered into OTUs at 97% identity and visualized through (A) relative abundance of Archaea, and (B) PCoA projection of the Weighted Unifrac dissimilarity matrix. Error bars represent standard deviation; significance bars represent group significance based on a two tail t-test, and stars denote the p-value thresholds (*=0.01, **=0.001, ***=0.0001).

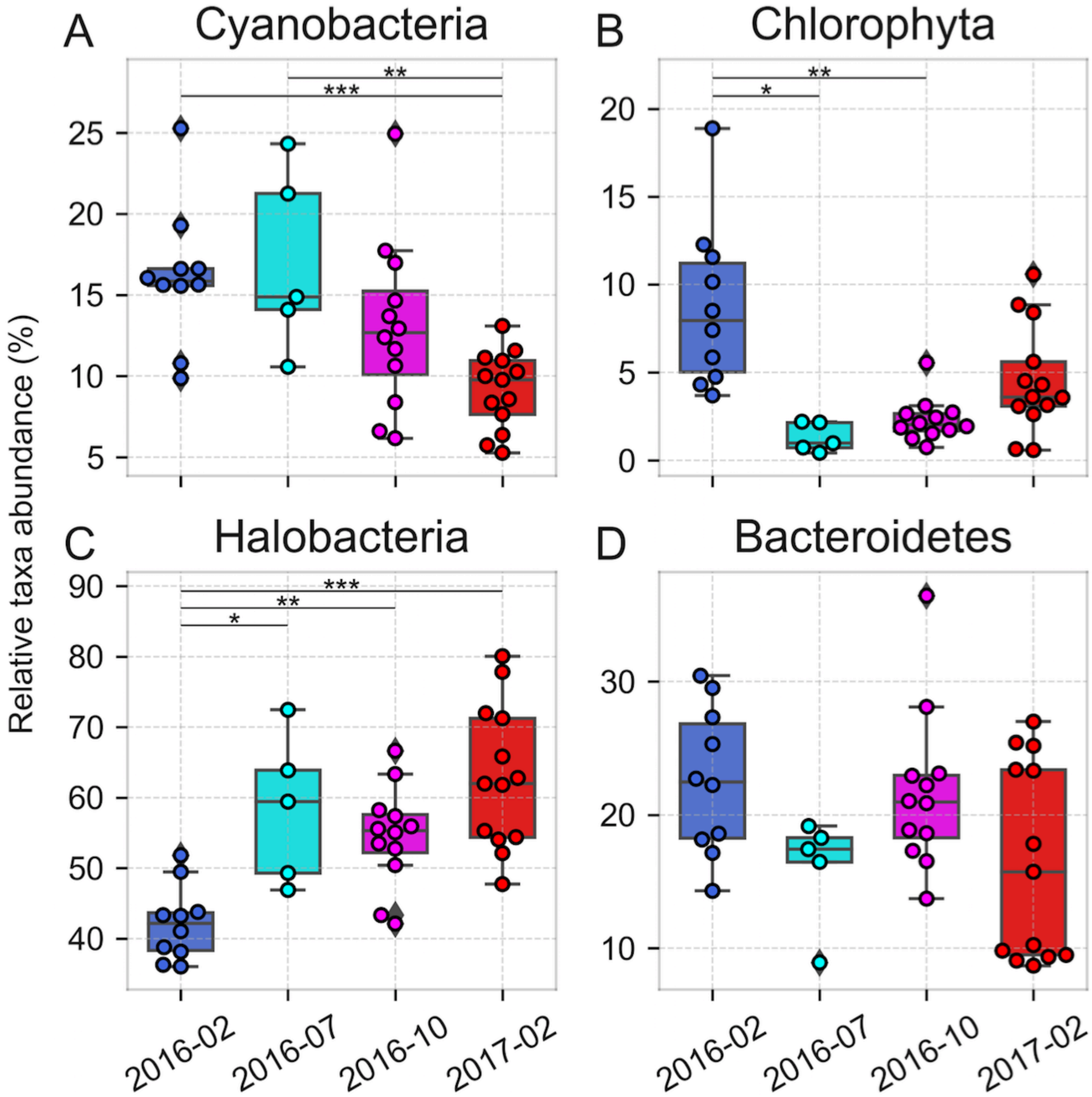


**Fig. S5.** Taxonomic composition of halite nodules harvested post-rain from Site 2 over time, inferred from 16S rRNA gene sequences clustered into OTUs at 97% identity and visualized through the relative abundance of dominant phyla (Chloroplast was used as a proxy for Chlorophyta and Halobacteria was the only class of Euyarchaeota) (A-D) Error bars represent standard deviation; significance bars represent group significance based on a two tail t-test, and stars denote the p-value thresholds (*=0.01, **=0.001, ***=0.0001).

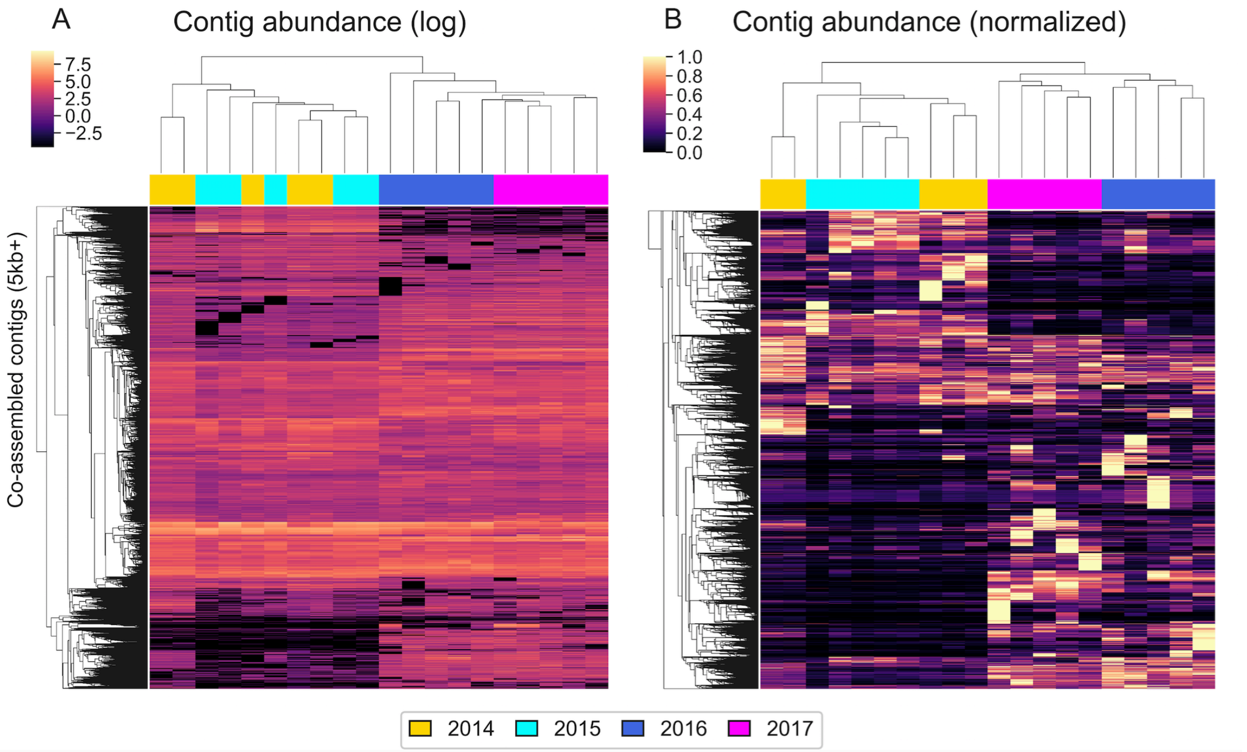


**Fig. S6.** Hierarchical clustering (Euclidean metric) of relative abundances (fragments per million) of contigs > 5kbp in the WMG co-assembly, quantified with reads from samples harvested at different dates and displayed on (A) a log scale and (B) standardized to the maximum abundance of each contig.

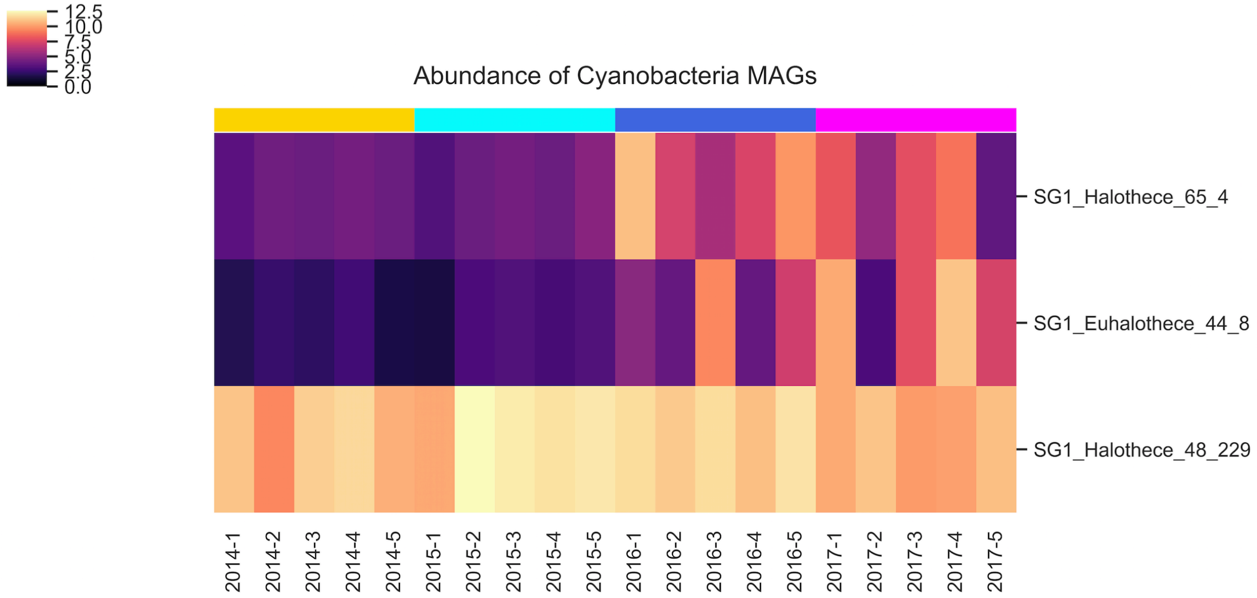


**Fig. S7.** Hierarchical clustering (Euclidean metric) of photosynthetic MAG relative abundances (fragments per million), quantified with metaWRAP's quant_bins module, showing the emergence of two new *Cyanobacteria* MAGs after the rain.

**Data S1.** Summary table of 16S rRNA gene OTUs clustered at 97% for Site 1 and Site 2, including OTU abundances across replicates, taxonomy, representative sequences, and stacked taxonomy plots visualizing community composition across the time-points and replicates.

**Data S2.** Summary table of reconstructed metagenome-assembled genomes (MAGs) with information about sequence statistics, binning accuracy estimated with CheckM, assembly coverage, taxonomy, and abundance across replicates in the time series.

**Tables (Supplementary):**

| Site | Latitude | Longitude | Elevation (asl) | Collection dates | Amplicon sequencing replicates | Shotgun sequencing replicates | Purpose |
|---|---|---|---|---|---|---|---|
| S1 | 20°57' 12.006"S | 70°1' 10.5996"W | 680m | Sep-14 | 10 | 5 | Before-after rain comparison |
| | | | | Jun-15 | 9 | 5 | |
| | | | | 8-Feb-16 | 19 | 5 | |
| | | | | 20-Feb-17 | 17 | 5 | |
| S2 | 20°57' 8.5212"S | 70°1' 1.2612"W | 664m | 8-Feb-16 | 12 | NA | After rain recovery process |
| | | | | 11-Jul-16 | 5 | NA | |
| | | | | 20-Oct-16 | 12 | NA | |
| | | | | 20-Feb-17 | 13 | NA | |
| S3 | 20°55' 48.18"S | 70°0' 49.32"W | 676m | Misc. | NA | 15 | Assembly and binning improveme nt |

**Table S1.** Description of sampling locations, dates, and replicate counts of biological samples collected for this study.
